# Microstructural Abnormalities of Substantia Nigra in Parkinson’s disease: A Neuromelanin Sensitive MRI Atlas Based Study

**DOI:** 10.1101/686154

**Authors:** Apoorva Safai, Shweta Prasad, Jitender Saini, Pramod Kumar Pal, Madhura Ingalhalikar

## Abstract

Microstructural changes associated with degeneration of dopaminergic neurons of the substantia nigra pars compacta (SNc) in Parkinson’s disease (PD) have been studied using Diffusion Tensor Imaging (DTI). However, these studies show inconsistent results, mainly due to methodological variations in delineation of SNc. To mitigate this, our work aims to construct a probabilistic atlas of SNc based on a Neuromelanin sensitive MRI (NMS-MRI) sequence and demonstrate its applicability to investigate microstructural changes on a large dataset of PD. Using manual segmentation and deformable registrations, we create a novel SNc atlas in the MNI space using NMS-MRI sequences of 27 healthy controls (HC). We employ this atlas to evaluate the diffusivity and anisotropy measures, derived from diffusion MRI in the SNc of 135 patients with PD and 99 HCs. Our observations of significantly increased diffusivity measures provide evidence of microstructural abnormalities in PD. However, no changes in the anisotropy were observed. Moreover, the asymmetry in abnormalities is prominent as the left SNc showed significant increase in diffusivity, and a reduction in FA when compared to the right SNc. Further the diffusivity and FA values also demonstrated a trend when correlated with the PD severity scores. Overall, from this work we establish a normative baseline for the SNc region of interest using NMS-MRI while the application on PD data emphasizes on the contribution of diffusivity measures rather than anisotropy of white matter in PD.

## Introduction

Parkinson’s disease (PD) is a chronic, progressive disorder characterized by bradykinesia, rigidity, and tremor. These symptoms have been predominantly implicated to degeneration of pigmented dopaminergic neurons in the A9 region of the substantia nigra pars compacta (SNc), which is a well-established early histological feature of PD (Fearnley & Lees, 1991). Degeneration of these nigral, dopaminergic neurons may lead to alterations in microstructural organization of the regional grey matter, white matter and local myelination of the SNc.

To extract the SNc accurately and gain deeper insight into the underlying volumetric and microstructural changes in this region of interest (ROI), conventional Magnetic Resonance Imaging (MRI) techniques have been used. For example, studies have employed T2 weighted, proton density weighted spin echo and single or dual inversion recovery based contrasts in identifying the SNc ROIs, with some reporting a significant difference in width, area or volume of SNc between PD patients and controls (Atasoy et al., 2004; Duguid, De La Paz, & DeGroot, 1986; Hutchinson & Raff, 2000; Pujol, Junque, Vendrell, Grau, & Capdevila, 1992; Stern, Braffman, Skolnick, Hurtig, & Grossman, 1989; Tuite, Mangia, & Michaeli, 2013), while others showed no significant difference (Geng, Li, & Zee, 2006; Oikawa, Sasaki, Tamakawa, Ehara, & Tohyama, 2002; Tuite et al., 2013). Recent studies have relied upon more sophisticated MRI protocols such as diffusion tensor imaging (DTI) either to ascertain the SNc ROIs (Menke, Jbabdi, Miller, Matthews, & Zarei, 2010; Sasaki et al., 2006) using fiber tractography or to evaluate the extent of this microstructural damage within the SNc in patients with PD. Table 1 provides a brief review of existing studies performed on the SNc, using diffusion MRI in PD. As observed from Table-1 there is a considerable variability in the reported findings that either illustrate no changes in FA(Aquino et al., 2014; Gattellaro et al., 2009; Menke et al., 2010; Schwarz et al., 2013), no changes in MD(Aquino et al., 2014; Du et al., 2011; Gattellaro et al., 2009; Menke et al., 2010; Peran et al., 2010; Rolheiser et al., 2011; Vaillancourt et al., 2009; Zhan et al., 2012), radial diffusivity(Du et al., 2011; Vaillancourt et al., 2009),or axial diffusivity(Du et al., 2011; Rolheiser et al., 2011; Vaillancourt et al., 2009), or demonstrate significant changes in these properties(Chan et al., 2007; Du et al., 2011; Knossalla et al., 2018; Langley et al., 2016; Loane et al., 2016; Peran et al., 2010; Rolheiser et al., 2011; Schwarz et al., 2013; Vaillancourt et al., 2009; Zhan et al., 2012). Moreover, in concurrence with the clinical asymmetry observed in PD, asymmetry in the FA values of the SNc have also been reported (Knossalla et al., 2018; Prakash, Sitoh, Tan, & Au, 2012). Taken together, the findings have been widely heterogeneous and perhaps could be implicated to inconsistencies in stages of disease severity between studies as well as to a relatively small sample size of these studies. More importantly, the techniques employed for delineating the SNc may have a significant bearing on the variability in the observed results. For instance, in a conventional T2 weighted MRI, there is variability in the hypointensity associated with the SNc as a result of increased iron deposition, in addition to the reduction in neuromelanin content(Deng, Wang, Yang, Li, & Yu, 2018; Langley et al., 2016; Wypijewska et al., 2010).This often leads to an inaccurate marking of SNc boundaries, false pathological representation of dopamine degeneration and also a poor contrast of the SNc(Langley et al., 2016). Diffusion tractography based techniques to localize the SNc are more complicated and may depend upon the choice of diffusion MRI protocol and the fiber tracking algorithm. Direct visualization and segmentation of the SNc, is therefore, a simpler yet a precise option to ensure superior accuracy in analysis of PD. To this end, a novel MR sequence known as “neuromelanin-sensitive MRI” (NMS-MRI) has demonstrated encouraging results. NMS-MRI which is a 3T T1 weighted high-resolution fast spin-echo sequence, is highly sensitive to the neuromelanin contained in the SNc and therefore renders the SNc as a hyperintense structure (Sasaki et al., 2008; Sasaki et al., 2006). This sequence is based on the paramagnetic properties of neuromelanin, a neuronal pigment which is a by-product of dopamine synthesis. Owing to the dopaminergic neuron loss in patients with PD, this normally hyperintense structure shows loss of normal signal intensity on NMS-MRI and therefore can be considered as a biomarker for PD. Multiple studies have demonstrated the clinical utility and accuracy of this sequence in patients with PD(Castellanos et al., 2015; Ohtsuka et al., 2014; Ohtsuka et al., 2013; Schwarz et al., 2011). To quantify these differences, the processing techniques have been based on visual inspection, or manual region of interest drawing followed by computation of volumes or contrast ratios(Isaias et al., 2016; Kashihara, Shinya, & Higaki, 2011; Matsuura et al., 2016; Matsuura et al., 2013; Ogisu et al., 2013; Ohtsuka et al., 2014; Reimao et al., 2015; Sasaki et al., 2006; Schwarz et al., 2011). Several T1 weighted NMS-MRI based studies in PD have found a reduction in signal intensity, contrast to noise ratio, as well as area and volume of SNc in PD patients as compared to controls (Isaias et al., 2016; Kashihara et al., 2011; Matsuura et al., 2016; Matsuura et al., 2013; Reimao et al., 2015; Sasaki et al., 2006). Majority of these studies have also showed a significant correlation of NMS-MRI measures with disease severity and other clinical scores(Isaias et al., 2016; Kashihara et al., 2011; Matsuura et al., 2016; Matsuura et al., 2013; Reimao et al., 2015), indicating a potential neuro-pathological biomarker for disease progression in PD.

**Table-1:**
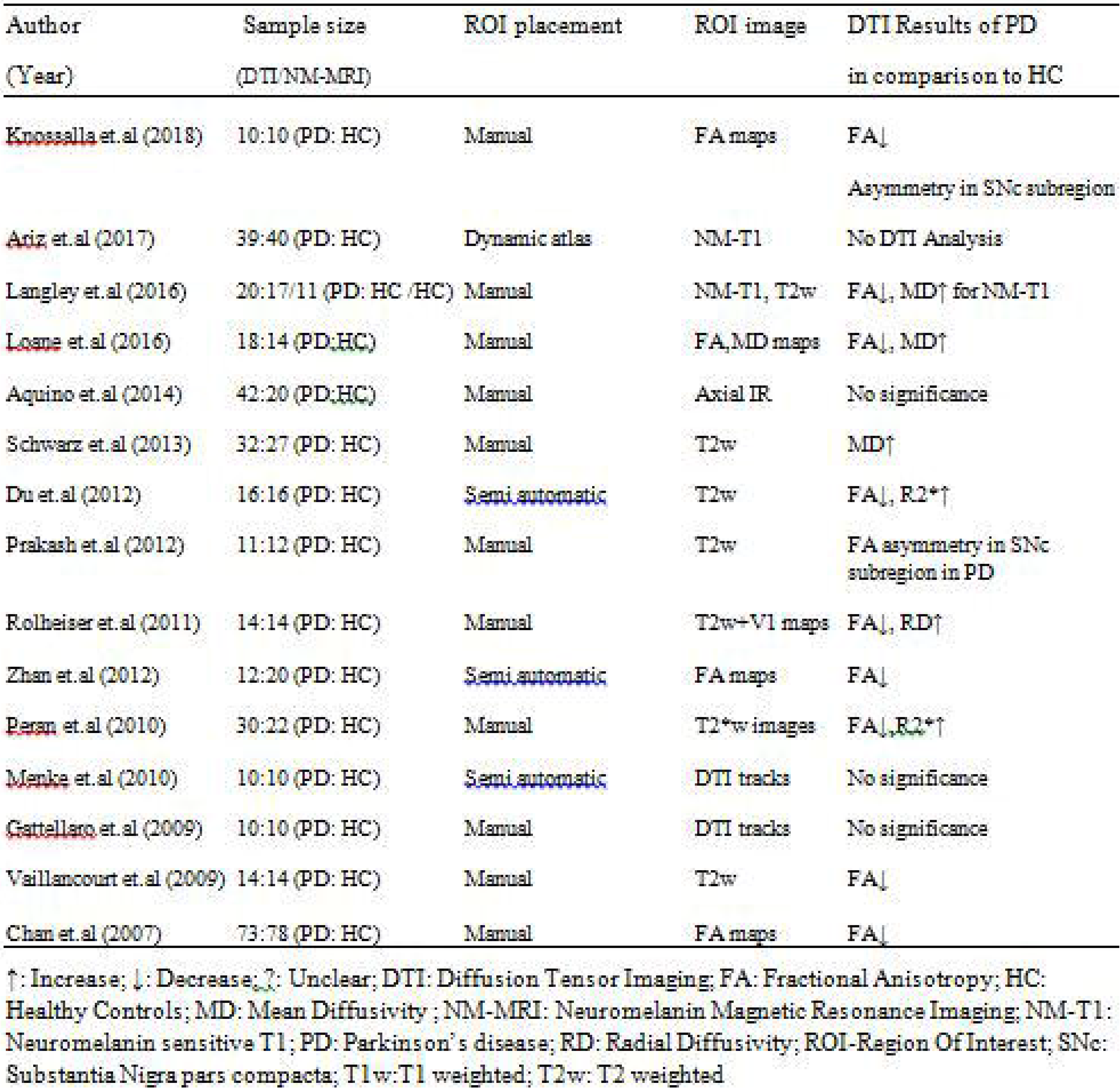
Review of articles that have performed Diffusion MRI analysis on the substantia nigra pars compacta in Parkinson’s disease

Considering the hyperintense SNc on NMS-MRI in healthy population, this sequence provides an opportunity to accurately localize and create a template of the SNc that can be utilized for analysis in parkinsonian disorders. Delineating the SNc in healthy controls and integrating these regions into a multi-image atlas creation process will not only offer better anatomical context to the template but also has the potential to facilitate superior segmentation. A recent study demonstrated creation of a dynamic SNc atlas, however, using a 2D NMS sequence (Ariz et al., 2018).

This study aims to generate a probabilistic atlas of the SNc from NMS-MRI and endeavors to apply the atlas on a large group of patients with PD to gain deeper insights into the diffusion MRI based microstructural abnormalities of the SNc. The generated probabilistic atlas of the SNc will provide a simple method for localization of the SNc, mitigating the methodological lack of uniformity in future studies.

## Methods

### Subject recruitment and clinical evaluation

One hundred and thirty-five subjects with PD and 99 healthy controls were recruited from two studies that were conducted at the Department of Neurology, National Institute of Mental Health and Neurosciences (NIMHANS), Bangalore, India. The diagnosis of idiopathic PD was based on the UK Parkinson’s Disease Society Brain Bank criteria(Hughes, Daniel, Kilford, & Lees, 1992) and confirmed by a trained movement disorder specialist (author PKP). Patients included in this study have been part of other studies(Lenka et al., 2018; Shah et al., 2017) from this group and all patients and controls provided informed consent prior to recruitment in the original projects.

Demographic and clinical details such as gender, age at presentation, age at onset of motor symptoms, disease duration, Mini Mental State Examination (MMSE), and Unified Parkinson’s Disease Rating Scale (UPDRS-III) OFF-state scores were recorded. Age and gender matched healthy controls with no family history of parkinsonism or other movement disorders were recruited.

Another group of 27 healthy controls whose NMS-MRI sequence was acquired as part of a different study (Prasad, Stezin, et al., 2018) of the same group, was used in the construction of our probabilistic atlas.

### Imaging

All subjects were scanned on a 3T Philips Achieva MRI scanner using a 32-channel head coil. Diffusion weighted images (DWI) for these subjects were acquired using a single-shot spin-echo EPI sequence with repetition time (TR)=8583-9070ms, echo time (TE)=60-62ms, field of view (FOV)=128×128×70m, voxel size=1.75×1.75×2mm, slice thickness=2mm. Diffusion gradient was applied in 15 directions, with b value =1000s/mm and a single b=0s/mm. T1-weighted images were acquired using TR=8.06ms, TE=3.6ms, voxel-size=1×1×1mm, FOV=256×256×160mm, slice thickness=1mm, voxel size=1×1×1mm and flip angle=8.

For creating a probabilistic atlas of SNc, a different set of controls were scanned on the same scanner. T1-weighted images were acquired using the above-mentioned protocol. Neuromelanin contrast sensitive sequences i.e. the spectral presaturation with inversion recovery (SPIR) sequence was acquired using TE: 2.2ms, TR: 26ms; flip angle: 20º; reconstructed matrix size: 512 × 512; field of view: 180×180×50mm; voxel size: 0.9×0.9×1mm; number of slices: 50; slice thickness: 1mm; and acquisition time: 4 minutes seconds. These images covered only the areas between the posterior commissure and inferior region of pons.

### Atlas construction

Probabilistic atlas was built from NMS-MRI images of 27 subjects as shown in Figure 1. For each subject, bilateral substantia nigra ROIs (snROIs) were created from NMS-MRI scans by manual segmentation. An author with expertise in the NMS-MRI sequence (SP), who was blinded to the groups, delineated the right and left SNc on the axial slices and created a 3D binary mask (snROIs). The NMS-MRI images were then linearly registered to the T1 image of the same subject by performing affine transformation using FLIRT in FSL(Smith et al., 2004). The T1 images of all subjects were pre-processed by performing motion correction, intensity inhomogeneity correction and skull stripping using Freesurfer 6.0(Fischl, 2012) and were transformed to the MNI space, by employing a deformable registration using Advanced Normalization Tools (ANTS), wherein a symmetric diffeomorphic transformation model (SyN) was applied and optimized using mutual information. The SyN is a large deformation registration algorithm which performs a bidirectional diffeomorphism and regularization using Gaussian smoothing of the velocity fields and has shown to outperform other nonlinear registration algorithms in preserving brain topology (Avants, Epstein, Grossman, & Gee, 2008).The transformations from NMS-MRI to T1 and from T1 to MNI were concatenated and were applied to the snROIs to transform them to MNI space. Along with visual inspection of each registered image, mean and variance of registered images was computed for quality check of registration. A probabilistic atlas of SNc in MNI space was then created, such that the voxel contained in all the 27 snROIs, was labelled with a probability of 1 and voxels not contained in any of the 27 ROIs were labelled with 0 probability.

**Figure-1:**
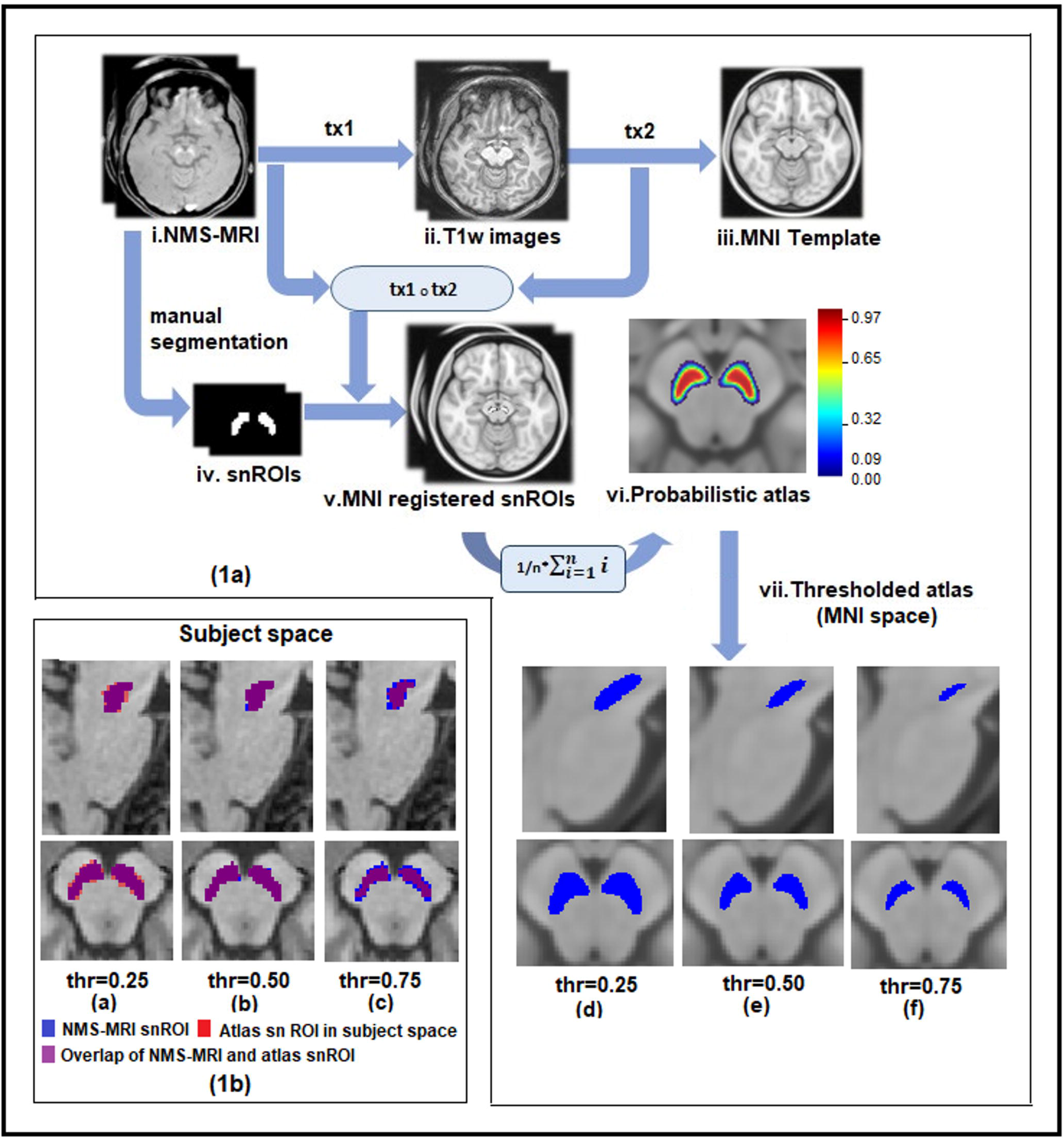
(a) Schematic for probabilistic and atlas construction of substantia nigra in MNI space with thresholding. i:i^th^ subject; n:number of subjects; tx1:transformation matrix of NMS-MRI to T1 registration, tx2:transformation matrix of T1 to MNI registration. (b) Probabilistic and thresholded substantia nigra atlas in subject space

### DTI pre-processing and analysis

DWI images of patients with PD healthy controls, were manually visualized for quality assessment. All pre-processing steps were done using FSL5.0.9 (Smith et al., 2004) which included removing the non-brain regions, correction for head movement and eddy current induced distortions using ‘eddy correct’ tool that performs an affine transformation between baseline b0 image and gradient images. The resulting rotating parameters of the affine transformation were used to rotate the gradients back, to align them with the transformed images. Least square approximation method was implemented to reconstruct the diffusion tensor images using ‘dtifit’, and the tensor fitting was checked for anatomical alignment. Diffusion maps such as fractional anisotropy (FA), mean diffusivity (MD), radial diffusivity (RD) and axial diffusivity (AD) were obtained by fitting the diffusion tensor model. FA maps of all subjects were registered to a standard FA map in MNI space-FMRIB58 image (https://fsl.fmrib.ox.ac.uk/fsl/fslwiki/FMRIB58_FA) using SyN algorithm and mutual information similarity index in ANTS. The transformation matrix of this registration was used to transform the MD, RD and AD maps to MNI space. Diffusion measures of bilateral SNc were extracted for all subjects using MNI registered diffusion maps and the atlas described in earlier, which was thresholded at 0.5 probability.

### Statistical Analysis

Statistical analysis on diffusion measures between PD and HC was performed using multivariate analysis of covariance (MANCOVA) model, with FA,MD,RD,AD of left and right SNc as dependant variables, PD and HC as grouping variables and age and gender as covariates. To evaluate associations between the microstructural changes to the severity of the disease, the residuals from the diffusion measures after regressing out age and gender were correlated to the UPDRS-III OFF scores (where available),the age of onset of disease (AoO), duration of illness (DoI) and levodopa equivalent daily dosage (LEDD) of patients with PD. To evaluate the asymmetry in diffusion measures of SNc, an independent t-test was done between age and gender corrected DTI measures of left and right SNc for PD patients and separately for healthy controls.

Further, to understand the discriminative power of these features, a multivariate classification framework with a random forest classifier was implemented to delineate PD from the healthy controls, with a 75%-25% train-test split on a total dataset of 234 subjects using all the diffusion measures of left and right SNc as the feature set. Classification performance was validated using a 10-fold cross validation (on the 75% training data) and evaluated using the test data. Relative feature importance was extracted to attain the contribution of each diffusion feature in classification of PD.

## Results

### Demographic and clinical details

Table-2 provides the complete demographic and clinical information for the dataset under consideration. There were no differences observed between the age and gender of the patient and control group. UPDRS-III OFF scores were available for 110 out of 135 subjects. With the exception of a single subject of PD, all others were right handed.

**Table-2:** Demographic and clinical characteristics of patients with Parkinson’s disease and healthy controls

|  | Parkinson's disease (n=135) | Healthy controls (n=99) |
| --- | --- | --- |
| Gender (F:M) | 66:69 | 60:39 |
| Age | 54.19 ± 8.23 | 52.34 ± 7.94 |
| Age at onset | 48.70 ± 9.36 | - |
| Duration of illness | 5.08 ± 3.11 | - |
| MMSE | 27.22 ± 2.51 | - |
| UPDRS III (OFF)* | 34.68 ± 8.80 | - |
| <u>Hoehn & Yahr</u> | 1.966 ± 0.611 | - |

F: Female; M: Male; MMSE: Mini mental status examination; UPDRS: Unified Parkinson's disease rating scale
\*: UPDRS III (OFF) was available for 110 subjects.

**Table-3:**
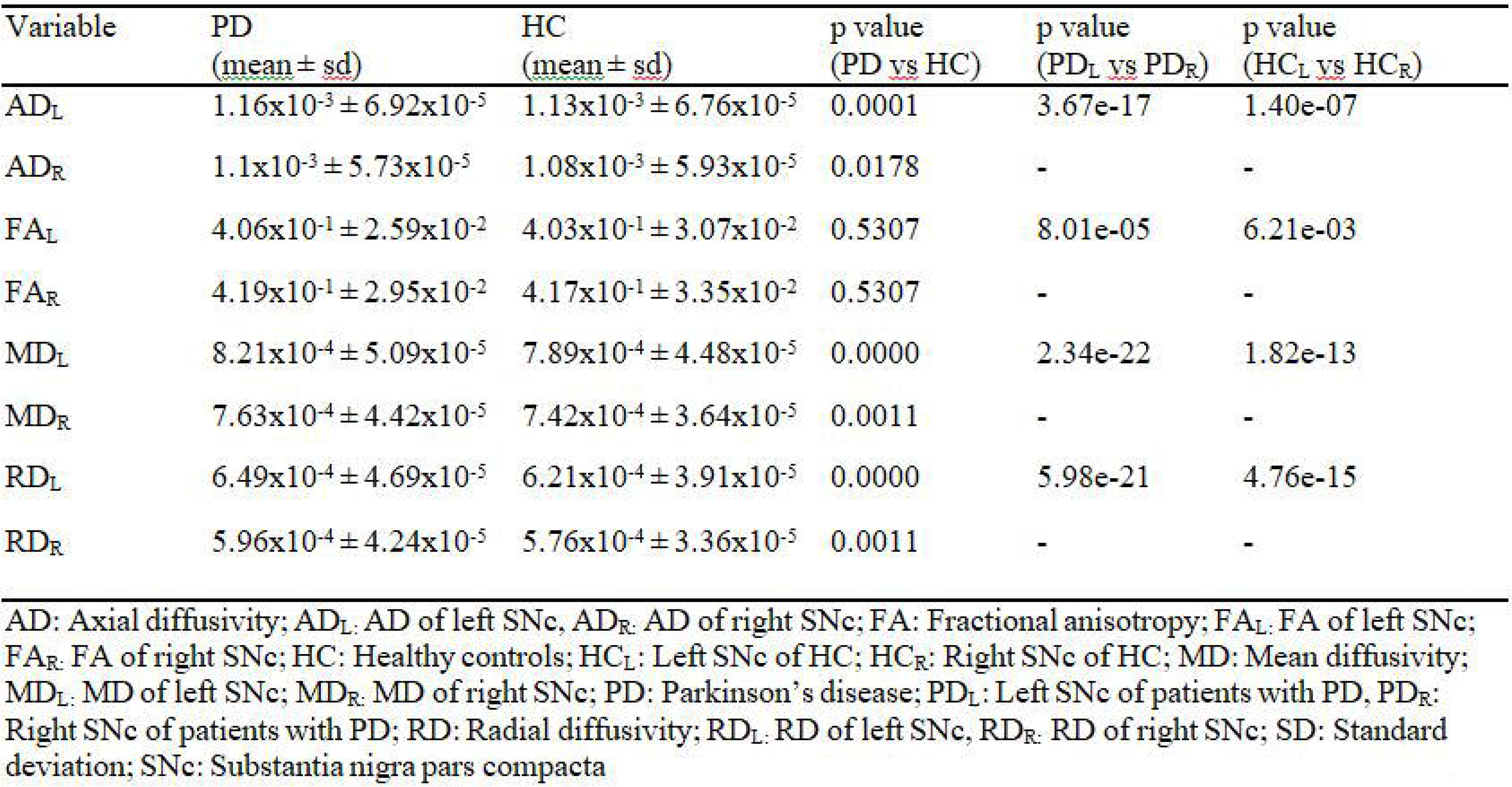
FDR corrected results of independent t-test for diffusion measures between healthy controls and patients with Parkinson’s disease

| Variable | PD<br>(<u>mean</u> ± <u>sd</u>) | HC<br>(<u>mean</u> ± <u>sd</u>) | p value<br>(PD <u>vs</u> HC) | p value<br>(PD<sub>L</sub> <u>vs</u> PD<sub>R</sub>) | p value<br>(HC<sub>L</sub> <u>vs</u> HC<sub>R</sub>) |
| --- | --- | --- | --- | --- | --- |
| AD<sub>L</sub> | 1.16x10<sup>-3</sup> ± 6.92x10<sup>-5</sup> | 1.13x10<sup>-3</sup> ± 6.76x10<sup>-5</sup> | 0.0001 | 3.67e-17 | 1.40e-07 |
| AD<sub>R</sub> | 1.1x10<sup>-3</sup> ± 5.73x10<sup>-5</sup> | 1.08x10<sup>-3</sup> ± 5.93x10<sup>-5</sup> | 0.0178 | - | - |
| FA<sub>L</sub> | 4.06x10<sup>-1</sup> ± 2.59x10<sup>-2</sup> | 4.03x10<sup>-1</sup> ± 3.07x10<sup>-2</sup> | 0.5307 | 8.01e-05 | 6.21e-03 |
| FA<sub>R</sub> | 4.19x10<sup>-1</sup> ± 2.95x10<sup>-2</sup> | 4.17x10<sup>-1</sup> ± 3.35x10<sup>-2</sup> | 0.5307 | - | - |
| MD<sub>L</sub> | 8.21x10<sup>-4</sup> ± 5.09x10<sup>-5</sup> | 7.89x10<sup>-4</sup> ± 4.48x10<sup>-5</sup> | 0.0000 | 2.34e-22 | 1.82e-13 |
| MD<sub>R</sub> | 7.63x10<sup>-4</sup> ± 4.42x10<sup>-5</sup> | 7.42x10<sup>-4</sup> ± 3.64x10<sup>-5</sup> | 0.0011 | - | - |
| RD<sub>L</sub> | 6.49x10<sup>-4</sup> ± 4.69x10<sup>-5</sup> | 6.21x10<sup>-4</sup> ± 3.91x10<sup>-5</sup> | 0.0000 | 5.98e-21 | 4.76e-15 |
| RD<sub>R</sub> | 5.96x10<sup>-4</sup> ± 4.24x10<sup>-5</sup> | 5.76x10<sup>-4</sup> ± 3.36x10<sup>-5</sup> | 0.0011 | - | - |

AD: Axial diffusivity; AD<sub>L</sub>: AD of left SNc, AD<sub>R</sub>: AD of right SNc; FA: Fractional anisotropy; FA<sub>L</sub>: FA of left SNc; FA<sub>R</sub>: FA of right SNc; HC: Healthy controls; HC<sub>L</sub>: Left SNc of HC; HC<sub>R</sub>: Right SNc of HC; MD: Mean diffusivity; MD<sub>L</sub>: MD of left SNc; MD<sub>R</sub>: MD of right SNc; PD: Parkinson's disease; PD<sub>L</sub>: Left SNc of patients with PD, PD<sub>R</sub>: Right SNc of patients with PD; RD: Radial diffusivity; RD<sub>L</sub>: RD of left SNc, RD<sub>R</sub>: RD of right SNc; SD: Standard deviation; SNc: Substantia nigra pars compacta

### Diffusion metrics

The probabilistic and thresholded (50%) atlas of the SNc computed from NMS-MRI images of 27 controls is shown in Figure-1. Significantly higher AD (left snROI: f=16.82,p-value<0.001; right snROI: f=6.5, p=0.011),MD(left snROI: f=21.40, p=0.000006; right snROI: f=11.43,p=0.00084) and RD(left snROI: f=19.57,p=0.000015; right snROI: f=11.77, p=0.0007) values were obtained in PD patients as compared to controls. As shown is Figure-2(a), the FA showed no significant differences (left snROI: f=0.511,p=0.476; right snROI: f=0.417. p=0.519). No significant correlations were obtained between clinical scores and diffusion measures; however, a correlation trend was observed between UPDRS and FA, duration of illness and FA, MD, RD and between FA and age of onset of disease as shown in supplementary figure-1. Asymmetry of all diffusion measures of bilateral SNc was observed for both HC and PD group, with PD group showing a higher significance. Left SNc showed higher AD(HC:t=5.521,p=1.07E-07;PD:t=9.085,p=2.36E-17),MD(HC:t=8.032,p=9.11E-14;PD:t=10.556,p=5.18E-22),RD(HC:t=8.724,p=1.22E15;PD:t=10.329,p=2.85E-21)as compared to right SNc. FA of the left SNc was found to be significantly lower as compared to right FA in the PD group(HC:t=-2.769,p=7.98E-05;PD:t=-4.007,p=7.98E-05).

Patients with PD were classified from HC with a 70.2% accuracy using random forest classifier, with MD and RD features showing higher contribution in the classification results as shown in Figure-2(b).

**Figure-2:**
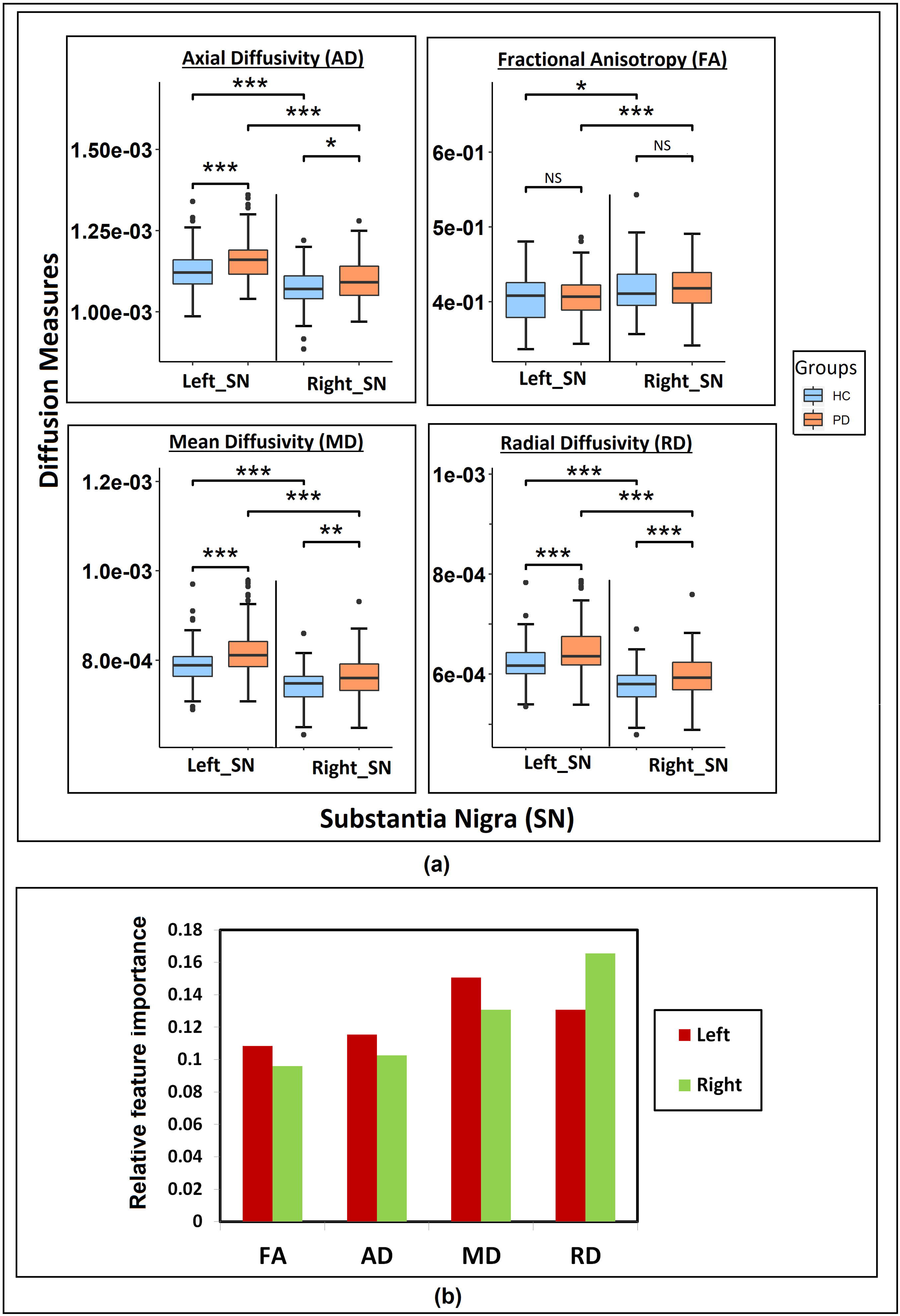
(a) Results of independent t-test indicating differences in diffusion measures of bilateral substantia nigra in Parkinson’s patient and healthy controls (b) Feature importance of DTI measures for classification of PD using random forest classifier.

## Discussion

We created a probabilistic atlas of the SNc by precisely extracting the SNc ROIs using NM rich MR sequence and employed it to accurately delineate the SNc to create a normative atlas that can be used in future PD studies. We applied this atlas to a large cohort of patients to PD to gain understanding of the microstructural abnormalities. Our results not only endorsed earlier findings but also facilitated fresh evidence supporting presence of micro-structural changes in PD substantia nigra compacta using diffusion MRI analysis. We demonstrated higher diffusivity values in the SNc in PD, with no changes in anisotropy and significant asymmetry of the diffusivity values.

SNc localization is particularly crucial to capture the subtle microstructural changes that occur as a consequence of dopaminergic neuronal loss in PD. Earlier DTI studies on PD have been based on contrast obtained from T2 weighted images(Du et al., 2011; Langley et al., 2016; Prakash et al., 2012; Rolheiser et al., 2011; Schwarz et al., 2013; Vaillancourt et al., 2009). Comparative studies(Du et al., 2011; Langley et al., 2015)on contrast generating MR sequence for SNc have shown that T2w sequences and NMS-MRI sequences are both sensitive to different sub regions of SNc, although hypo-density observed on T2 images in PD could also indicate iron deposition apart from dopaminergic loss in the SNc, making it highly unreliable(Deng et al., 2018; Langley et al., 2016; Wypijewska et al., 2010). Recent work by Visser et al.(Visser et al., 2016) has employed the FLASH sequence on 7T MRI to delineate the substantia nigra. However, this sequence does not capture the SNc, as it is sensitive only to the elevated concentrations of ferritin that are prominently observed in substantia nigra pars reticulata (SNr), as has been demonstrated by an earlier study (Langley et al., 2015). Therefore, applying the SNr atlas to PD is unreliable in understanding the abnormalities which occur predominantly in the SNc owing to the dopaminergic neuronal loss in PD. To alleviate these limitations, NMS-MRI has been employed to visualize and quantify the intensity contrast in SNc(Isaias et al., 2016; Matsuura et al., 2016; Matsuura et al., 2013; Ohtsuka et al., 2014; Ohtsuka et al., 2013; Reimao et al., 2015; Sasaki et al., 2006; Schwarz et al., 2011). The studies employing this contrast have corroborated its utility not only to render the SNc region but also as a contrast ratio-based biomarker in PD. However, no studies have comprehensively worked on creating a SNc template which is crucial to overcome the discrepancies in SNc localization, providing a normative baseline for comparison of results across studies. Our work constructs an accurate probabilistic atlas that will facilitate valid NM based SNc regions and thus ensures reliable results with DTI measures.

Our atlas creation is based on uniform and accurate image registration of all subjects to the MNI space. A review study on 14 different non-linear registration algorithms have found that ART and SyN algorithms have consistently performed well across multiple datasets(Klein et al., 2009). We have employed a symmetric diffeomorphic (SyN) registration using the ANTs toolbox for registering subject T1 images and subsequently the SNc masks, created from N M rich sequences of 27 subjects onto the MNI space as shown in Figure-1.The maximised optimization of space-time deformation maps in SyN and hierarchical interpolation performed in ANTs, increases normalization accuracy and preserves the brain topology, thus enhancing the registration precision of our probabilistic atlas(Avants et al., 2008; Klein et al., 2009). Each of the registrations was manually checked for precision in registration. The probabilistic atlas created, was thresholded at 50% probability, as it removed the voxels outside the expected SNc region as shown in Figure 1.

Our analysis of diffusion measures was performed on a large cohort of patients with PD, for the first time, where we observed a significantly increased MD, RD and AD in the patients compared to age and gender matched healthy controls (Figure-2(a)), with no significant differences in FA. Earlier studies have reported mixed results for significant differences in FA as shown in Table-1. Although our results appear to contradict these studies, the current study has a higher statistical power and perhaps a more robust SNc region of interest, making it more relevant. The significant changes were obtained only in the diffusivity measures (MD,RD,AD) and were also reflected in the classifier results where the left and right MD and RD were captured as the most discriminative features of PD as shown in Figure 2(b). Moreover, these microstructural features of diffusivity could classify patients with PD from HC with an accuracy of 70.2%, thereby facilitating a novel biomarker for PD.

Degree of myelination, axonal diameter and distance between extracellular membranes drive the changes in radial diffusivity, whereas diffusion anisotropy implies a directional alignment of white matter tracts(Beaulieu, 2002).Intuitively, the biological process of fibre disintegration and de-myelination which are associated with neuro degeneration, should lead to an increase in RD and reduction in FA values. However, neuro-degeneration may involve multiple additional pathological processes such as changes in membrane permeability, re-structuring of white matter fibres, glial alterations and damage to the intracellular compartment. The degree of variation in these processes may be contributing towards the proportional changes in diffusion tensors in all three dimensions, and thereby reducing the sensitivity of FA(Acosta-Cabronero, Williams, Pengas, & Nestor, 2010). Nevertheless, it is important to note that our diffusion protocol was limited to 15 gradient directions which may not facilitate the best model (Jones, Knosche, & Turner, 2013) for fiber tractography or connectivity, but is valid for computing diffusivity and anisotropy measures.

In addition to this, our findings on asymmetry of diffusion measures in SNc support the speculated theory of asymmetrical degeneration of dopaminergic nigral neurons in PD. Even though we do not capture any significant differences in FA, we observe that the FA in left SNc reduces significantly when compared to the FA in right SNc, with higher significance in patients with PD in comparison to HC(Figure-2). Similarly, MD, AD and RD demonstrated significantly higher values in the left SNc when compared to right (Figure-2). Earlier work by our group has illustrated such asymmetry using contrast ratios on NMS-MRI in the SNc(Prasad, Saini, Yadav, & Pal, 2018). Our correlation analysis did not demonstrate any significant associations of DTI measures with AoO, DoI, UPDRS-III OFF, or LEDD scores. However, a trend was observed between UPDRS, and FA, DoI and FA, MD, RD and between AoI and FA (Supplementary figure-1).

## Conclusions

In conclusion, this study addressed a crucial question of uniform SNc localization in patients with PD and performed a large-scale robust assessment of microstructural features of the SNc in PD using diffusion measures. Our standardised SNc atlas based on the NMS-MRI will be released to the scientific community and will further aid in eliminating the methodological variability associated with delineation of SNc. Microstructural abnormalities of the SNc in PD are predominantly associated with altered diffusion metrics rather than anisotropy, and demonstrate a significant asymmetry which is in concurrence with clinical lateralization of symptoms. These results obtained from our large-scale study on an accurately delineated SNc provide a thorough and reliable profile of the neuro-degeneration associated microstructural abnormalities of the SNc in PD.

## Acknowledgements

This work was financially supported by DST SERB (ECR/2016/000808), Indian Council of Medical Research (Grant Number ICMR/003/304/2013/00694) and Department of Biotechnology (Grant Number: BT/PR14315/MED/30/474/201

**Supplementary Figure-1:** Correlations between DTI measures and clinical scores which showed borderline significance.

