## Supplementary_material for "Microstructural Abnormalities of Substantia Nigra in Parkinson’s disease: A Neuromelanin Sensitive MRI Atlas Based Study"

Correlation Analysis:

Pearson's correlation was performed on all DTI measures and age and gender regressed, residual clinical scores such as UPDRS III (off), Age of Onset (AoI),Duration of Illness (DoI) and Levodopa Equivalent Daily Dosage (LEDD).Although no signficant correlations were obtained, a trend was observed between diffusion measures of left SNc and clinical scores. FA of left SNc showed a negative correlation trend with DoI (r=-0.154, p=0.091) and UPDRS scores (r=-0.180, p=0.06),and a positive trend with AoI (r=0.14, p=0.104). MD (r=0.1, p=0.273) and RD (r=0.13, p=0.154) both showed a positive correlation trend with DoI.

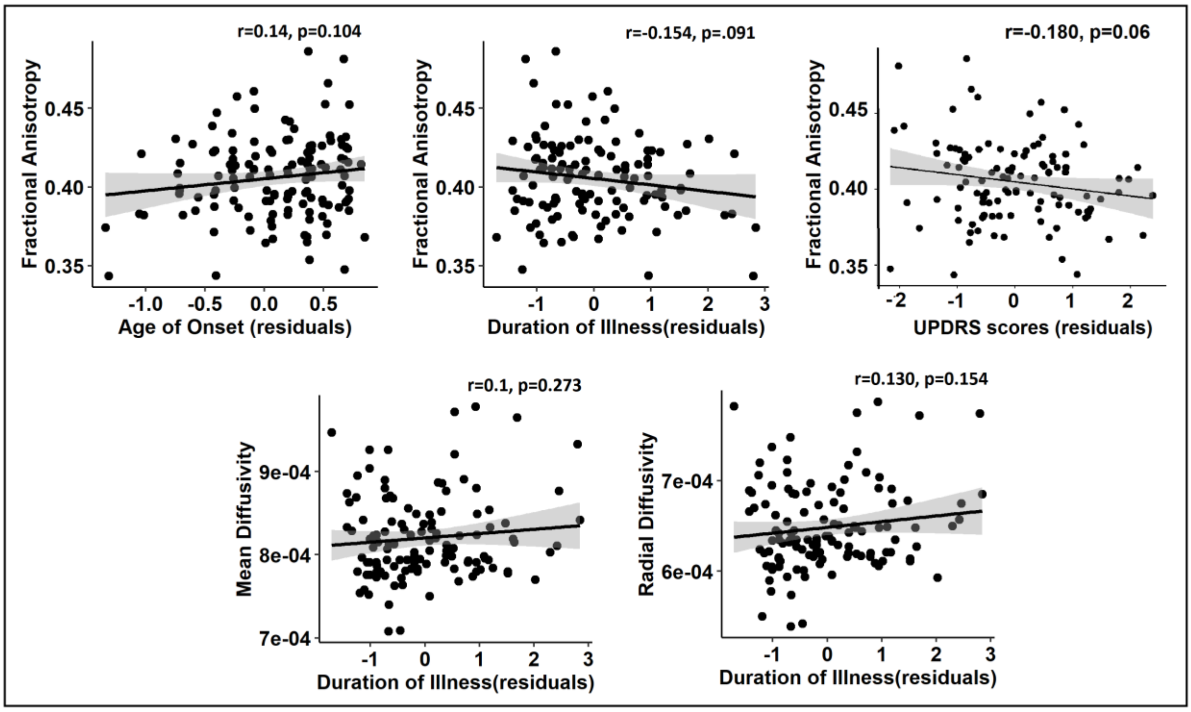


Figure 1: Correlations between DTI measures of left SNc and residual clinical scores with age and gender regressed.
